# Fibrillin 2 interacts with other proteins to protect photosystem II against abiotic stress in *Arabidopsis thaliana*

**DOI:** 10.1101/2020.10.07.329979

**Authors:** Diego Torres-Romero, Ángeles Gómez-Zambrano, Antonio Jesús Serrato, Mariam Sahrawy, Ángel Mérida

**Affiliations:** Instituto de Bioquímica Vegetal y Fotosíntesis. Consejo Superior de Investigaciones Científicas (CSIC) – Universidad de Sevilla. Avenida Américo Vespucio, 49. 41092-Sevilla. Spain; Estación Experimental del Zaidín. Consejo Superior de Investigaciones Científicas (CSIC). Calle Profesor Albareda, 1. 18008-Granada. Spain

## Abstract

Fibrillins (FBNs) are plastidial proteins found in photosynthetic organisms, from cyanobacteria to higher plants. The function of most FBNs is largely unknown. We focused on the subgroup formed by FBN1a, −1b, and −;2, which has been proposed to be involved in the photoprotection of photosystem II (PSII), though their mechanism of action has not yet been characterized. We show that FBN2 interacts with FBN1a and with other FBN2 polypeptides, potentially forming a network around the plastoglobule surface. Both FBN2 and FBN1 interact with the allene oxide synthase, and the elimination of any of these FBNs results in a delay in jasmonate-mediated anthocyanin accumulation in response to a combination of moderate-high light and low temperature. FBN2 also interacts with other proteins involved in different metabolic processes. Mutants lacking FBN2 demonstrate less photoprotection of PSII, alterations that are not found in *fbn1a-fbn1b* mutants. We also show that FBN2 interacts with Acclimation of Photosynthesis to Environment 1 (APE1), and gene co-expression analysis suggests that both proteins are involved in the same metabolic process. The elimination of APE1 leads to lesions in PSII under abiotic stress similar to observations in *fbn2* mutants, with lower maximum and effective quantum yield. However, a reduction in non-photochemical quenching is observed exclusively in *fbn2* mutants, suggesting that other FBN2-interacting proteins are responsible for this alteration. We propose that FBN2 facilitates accurate positioning of different proteins involved in distinct metabolic processes, and its elimination leads to dysfunction in those proteins.

**One sentence summary:** Fibrillin 2 protects photosynthesis against abiotic stresses by facilitating the accurate positioning of different proteins involved in distinct processes, and its elimination leads to dysfunction in those proteins

## INTRODUCTION

Plastoglobules (PGs) are lipoprotein bodies present in all types of plant plastids. In chloroplasts, they are attached to thylakoids through a half-lipid bilayer that surrounds the globe contents and is continuous with the stroma-side leaflet of the thylakoid membrane (Austin et al., 2006). The lipids of chloroplast PGs consist mainly of prenylquinones and triacylglycerol, and PGs have been proposed to serve as lipid microcompartments for the synthesis, storage, and redistribution of these subsets of isoprenoids and neutral lipids (van Wijk and Kessler, 2017). However, proteomic analysis has shown the presence of different proteins in PGs (Vidi et al., 2006; Ytterberg et al., 2006). Lundquist et al. (2012) defined a PG proteome core consisting of 30 proteins, which were classified into four modules using a genome-wide co-expression analysis: Module 1 (senescence, chlorophyll degradation, proteolysis), Module 2 (plastid carotenoid metabolism, plastid proteolysis), Module 3 (redox regulation, photoacclimation), and Module 4 (plastid biogenesis and Calvin-Benson cycle). These studies demonstrated that PGs are not a simple lipid reservoir, and proteins involved in different metabolic processes of the chloroplast are associated with them. The most abundant proteins in PGs are the fibrillins (FBNs) and six ABC1 kinases, constituting 53% and 19% of the core PG proteome mass, respectively (Lundquist et al., 2012). The FBNs are a large protein family present in photosynthetic organisms, from cyanobacteria to higher plants (Laizet et al., 2004). The FBNs in higher plants and algae can be divided into 12 clades (Singh and McNellis, 2011). Seven of them are located in PGs and form part of its proteome core (FBN1a, −1b, −2, −4, −7a, −7b, and −8), and the rest have a stromal or thylakoid-associated location (Lundquist et al., 2012). FBNs have been proposed to be involved in central processes of plant physiology, particularly plant responses to stress and plastid architectural development (Singh and McNellis, 2011). However, their mechanisms of action have not yet been elucidated. Recently, stromal localized FBN5 was shown to be essential for plastoquinone-9 biosynthesis by binding to solanesyl diphosphate synthases in *Arabidopsis* (Kim et al., 2015), and the elimination of FBN6 leads to perturbation of ROS homeostasis (Lee et al., 2019). FBN1a and FBN1b arose from a recent duplication event on chromosome 4, and together with FBN2 form a subgroup within the FBN family. Most of the FBNs characterized to date in different species belong to this subfamily (Laizet et al., 2004) and, together with FBN4, are the most abundant proteins in PGs (Lundquist et al., 2012). The function of the subgroup formed by FBN1a, −1b, and −2 has been analyzed using an RNA interference strategy that reduces the expression of these three genes (Youssef et al., 2010). This reduction leads to pleiotropic alterations, such as abnormal granal and stromal membrane arrangement, higher photosystem II (PSII) photoinhibition under stress, retarded shoot growth, and a deficit in anthocyanin accumulation under stress. These phenotypic alterations were abolished by jasmonate (JA) treatment, and light/cold stress-related JA biosynthesis has been suggested to be conditioned by the accumulation of PG-associated FBN1-2 proteins (Youssef et al., 2010), though the mechanism explaining the function of the FBNs is not clear.

In this study, we dissect the function of the FBN1-2 subgroup by analyzing the phenotype of *fbn2*, *fbn1a-fbn1b*, and *fbn1a-1b-2* knock-out mutant lines. We show that FBN2 interacts with FBN1a, with another FBN2 polypeptide, and with other proteins, including the enzymes catalyzing the first steps of the synthesis of JA. The interaction seems to be necessary for the correct function of these proteins. FBN1a, −1b, and −2 operate together in some processes but independently in others, such as protection of photosynthesis performance under light and temperature stress. This protection seems to be exerted by interaction of FBN2 with proteins, such as Acclimation of Photosynthesis to Environment (APE1) and other yet uncharacterized proteins.

## RESULTS

### FBN2 is localized in both the stroma and in association with membranes

Both FBN1a and FBN1b exhibit a dot-like pattern of localization when analyzed using transient expression of FBN1-GFP fusion peptide in *Nicotiana* leaves, which is characteristic of proteins associated with PGs (Gámez-Arjona et al., 2014a). However, FBN2-GFP exhibited a different pattern, and the detected fluorescence coincided with the chlorophyll autofluorescence (Figure 1A), indicating that FBN2 could be a soluble protein of the stroma or evenly associated with thylakoid membranes. Immunoblot analysis of the soluble and membrane fractions from isolated chloroplasts indicated that FBN2 has dual localization, both soluble in the chloroplast stroma and associated with membranes (Figure 1B). With this analysis, we cannot ascertain whether the FBN2 population associated with membranes is localized exclusively in association with PGs or can also be associated with other regions of the thylakoid membranes.

**Figure 1.**
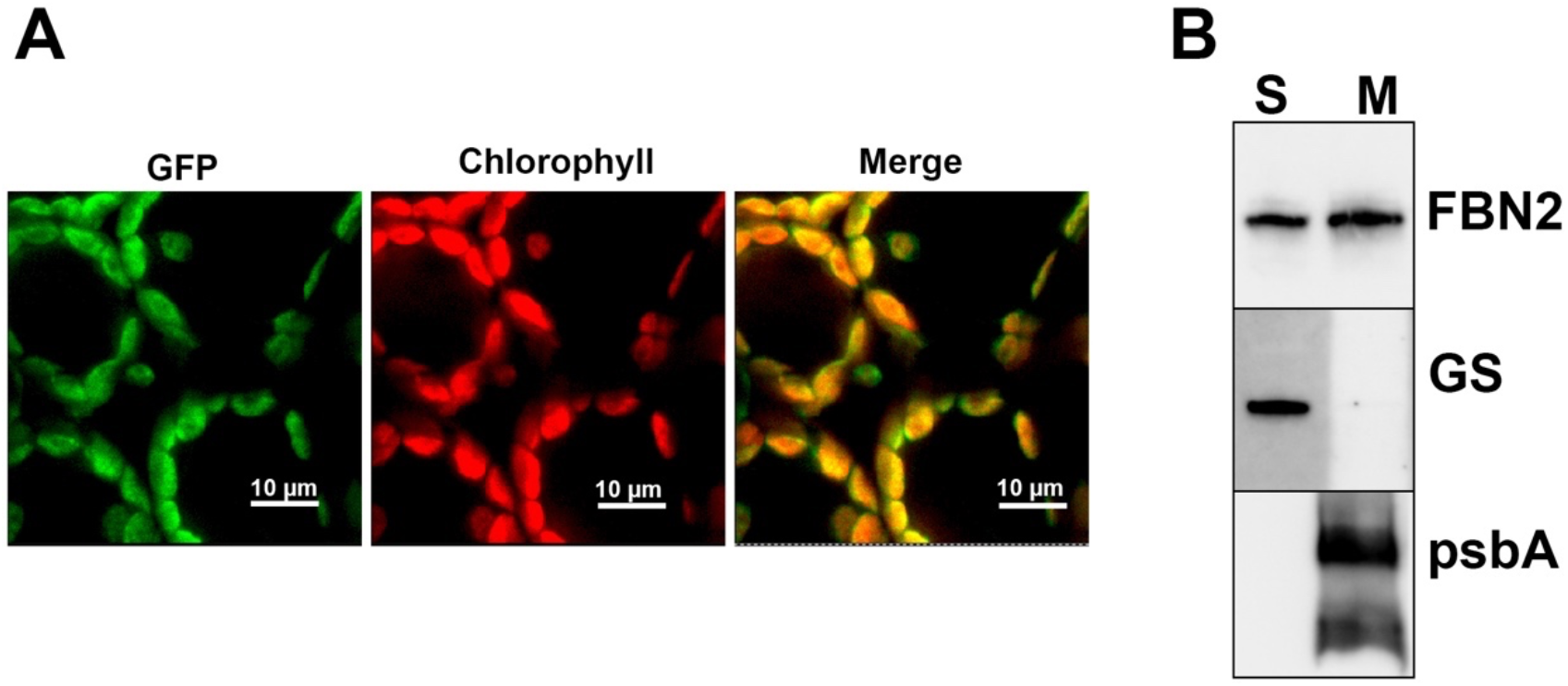
Localization of FBN2. (A) Full-length FBN2 cDNA was fused to GFP and transiently expressed in *Nicotiana benthamiana* leaves. Fluorescence was monitored by confocal microscopy. The GFP fluorescence, chlorophyll autofluorescence, and merged images are shown. (B) Chloroplasts isolated from *Arabidopsis* rosette leaves were disrupted and the soluble (S) and membrane fractions (M) isolated by ultracentrifugation at 100,000 × g for 1 hour at 4°C. The pellet (membrane fraction) was resuspended in the same volume as the supernatant and 20 μl of each fraction used in SDS-PAGE. Proteins were blotted onto a PVDF filter and hybridized with specific antibodies against FBN2, plastidial glutamine synthetase (GS) as a marker of stroma protein, and psbA as marker of thylakoid membranes.

### Effect of cold and high-light stress on growth and anthocyanin accumulation in mutants affected in FBN1a, −1b, and −2

Previous studies have shown that *fbn1a-1b-2* knock-down mutant plants have reduced growth and accumulate lower levels of anthocyanins than wild-type (WT) plants when subjected to cold and high-light stress (Youssef et al., 2010). To dissect the role of each protein in these phenotypic alterations, we analyzed the growth rate and anthocyanin accumulation in leaves of knock-out mutants lacking FBN2 (*fbn2*), FBN1a and FBN1b (*fbn1a-1b*) or FBN2, FBN1a and FBN1b (*fbn1a-1b-2*). We did not analyze the single knock-out mutants *fbn1a* and *fbn1b* because these proteins have 89% identity in the amino acid sequence of the mature proteins, and we have shown previously that these polypeptides can form homodimers and heterodimers (Gámez-Arjona et al., 2014a), suggesting a high degree of functional redundancy between these two proteins. In accordance with previous data, we did not detect altered growth in *fbn2*, *fbn1a-1b,* and *fbn1a-1b-2* plants cultivated under normal growth conditions (110 μmol m^−2^ s^−1^ light intensity, 22°C day/20°C night temperature). However, in contrast to what has been described for *fbn1a-1b-2* knock-down plants, we did not find any growth retardation when plants were cultivated under normal conditions over 19 days and then subjected to cold and high light stress (600 μmol m^−2^ s^−1^ light intensity, 10°C temperature) over 30 days (Figure 2A). The RNA interference strategy employed in previous studies (Youssef et al., 2010) could affect the levels of expression of other FBN genes and may explain the discordance with our results. High light induced the accumulation of anthocyanins in the different lines. The induction was similar in the three mutant lines and considerably lower than the induction measured in WT plants after 1 week of stress (Figure 2B). The reduced accumulation of anthocyanins in leaves was similar in the *fbn2*, *fbn1a-1b*, and *fbn1a-1b-2* lines, suggesting that FBN2 and FBN1s act together to facilitate the accumulation of anthocyanins. Nevertheless, the levels of anthocyanins in the leaves of the different mutants reached the values detected in WT plants after 3 weeks of stress (Figure 2B), indicating that the accumulation of anthocyanins was impaired, but not abolished, in the mutant lines. The delay in the accumulation of anthocyanins was abolished by treatment with JA (Figure 2C), supporting the idea that this delay is a consequence of impaired synthesis of JA in the mutants.

**Figure 2.**
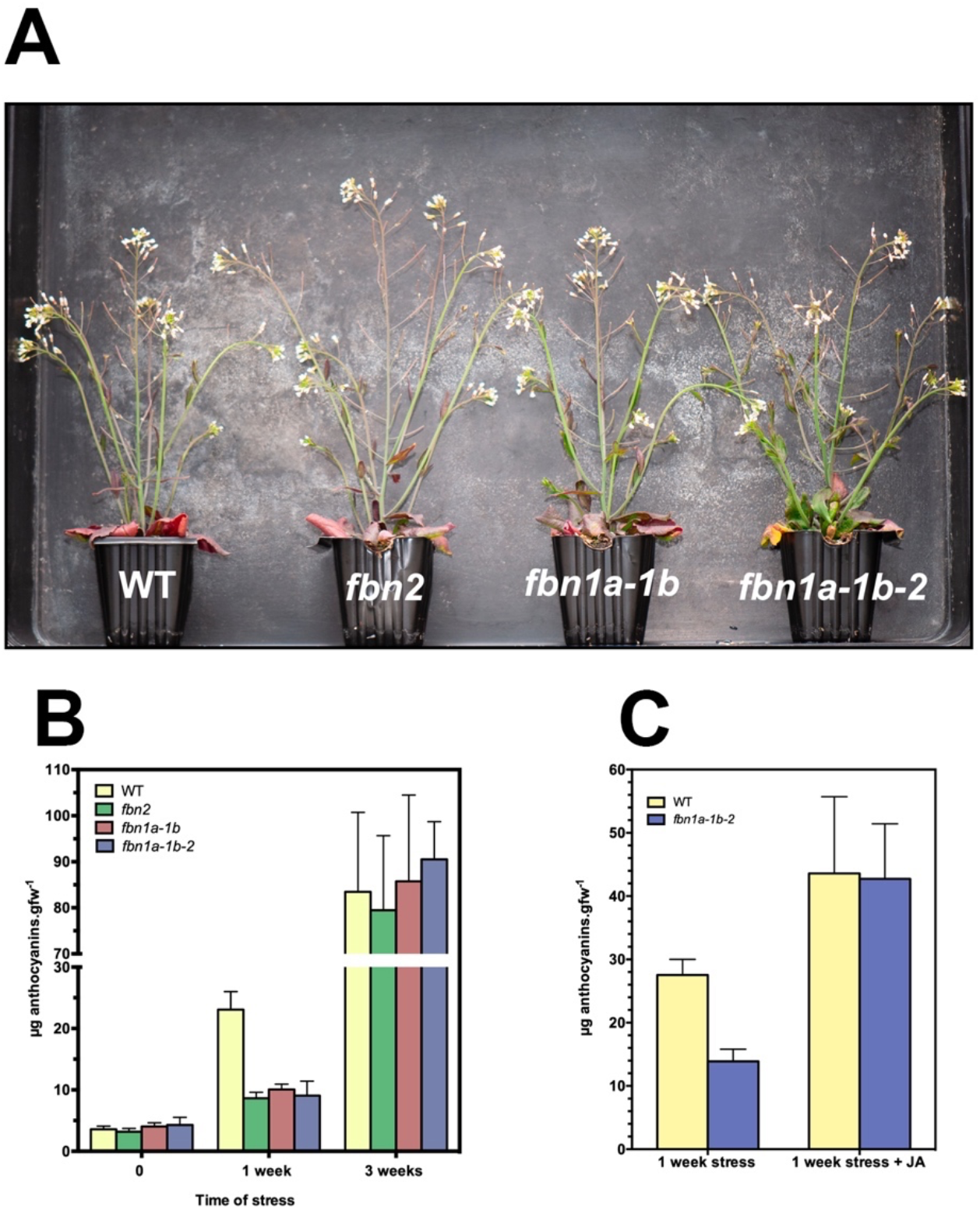
Phenotypic alterations in *fbn1* and *fbn2* mutants. (A) Knock-out mutants *fbn2*, *fbn1a-1b,* and *fbn1a-1b-2*, and WT plants were sown in soil and cultivated under normal conditions for 3 weeks. After this period, plants were transferred to stress conditions (600 μmoles m^−2^ s^−1^ light intensity and 10°C temperature) for another 3 weeks and the growth of the plants documented. (B) Anthocyanin accumulation in *fbn* mutants. Leaves of plants grown and stressed as in (A) were harvested at time 0 and 1 and 3 weeks of treatment and the levels of anthocyanins determined. (C) Plants cultured under normal conditions were subjected to stress as described in (A) for 1 week with or without the addition of 2 mM jasmonate every 3 days and the levels of anthocyanins determined. Values are presented as the mean ± SD (n = 3).

### Effect of mutations on photosynthetic performance

One of the functions proposed for FBNs has been protection of the photosynthetic apparatus against damage promoted by different abiotic stresses (Singh and McNellis, 2011). We analyzed the performance of PSII by determining the chlorophyll a fluorescence of the different mutant plants subjected to abiotic stresses, such as high light, cold, or a combination of both. Under normal growth conditions, the values of the photosynthetic parameters studied in the mutant plants were indistinguishable from those in WT plants (Figure 3A and 3B). High light or cold stress alone did not affect these values (Figure S1A and S1B). However, the combination of high light and cold altered the studied parameters in the *fbn2* and *fbn1a-1b-2* mutants, but not *fbn1a-1b* mutant plants (Figure 3A and 3C). The elimination of FBN2 diminished the maximum quantum yield of PSII (F_v_/F_m_) with respect to WT plants after 24 hours of stress. This reduction was maintained after 1 week of stress (Figure 3A). It also decreased the effective PSII quantum yield (Φ_II_) and the yield of non-photochemical-regulated dissipation of energy (Φ_NPQ_); therefore, the yield of the non-regulated dissipation of energy (Φ_NO_) increased. Note that the sum of all yields for the dissipative processes of the energy absorbed by PSII is unity (Φ_NPQ_+Φ_II_+Φ_NO_=1) (Kramer et al., 2004). In addition, the excitation pressure (1-qL), the percentage of primary PSII electron acceptor that was reduced, increased in *fbn2* and *fbn1a-1b-2* mutants, but not in the double mutant *fbn1a-1b* (Figure 3C). These data point to a specific role of FBN2 in the protection of PSII against stresses, differentiating it from FBN1s.

**Figure 3.**
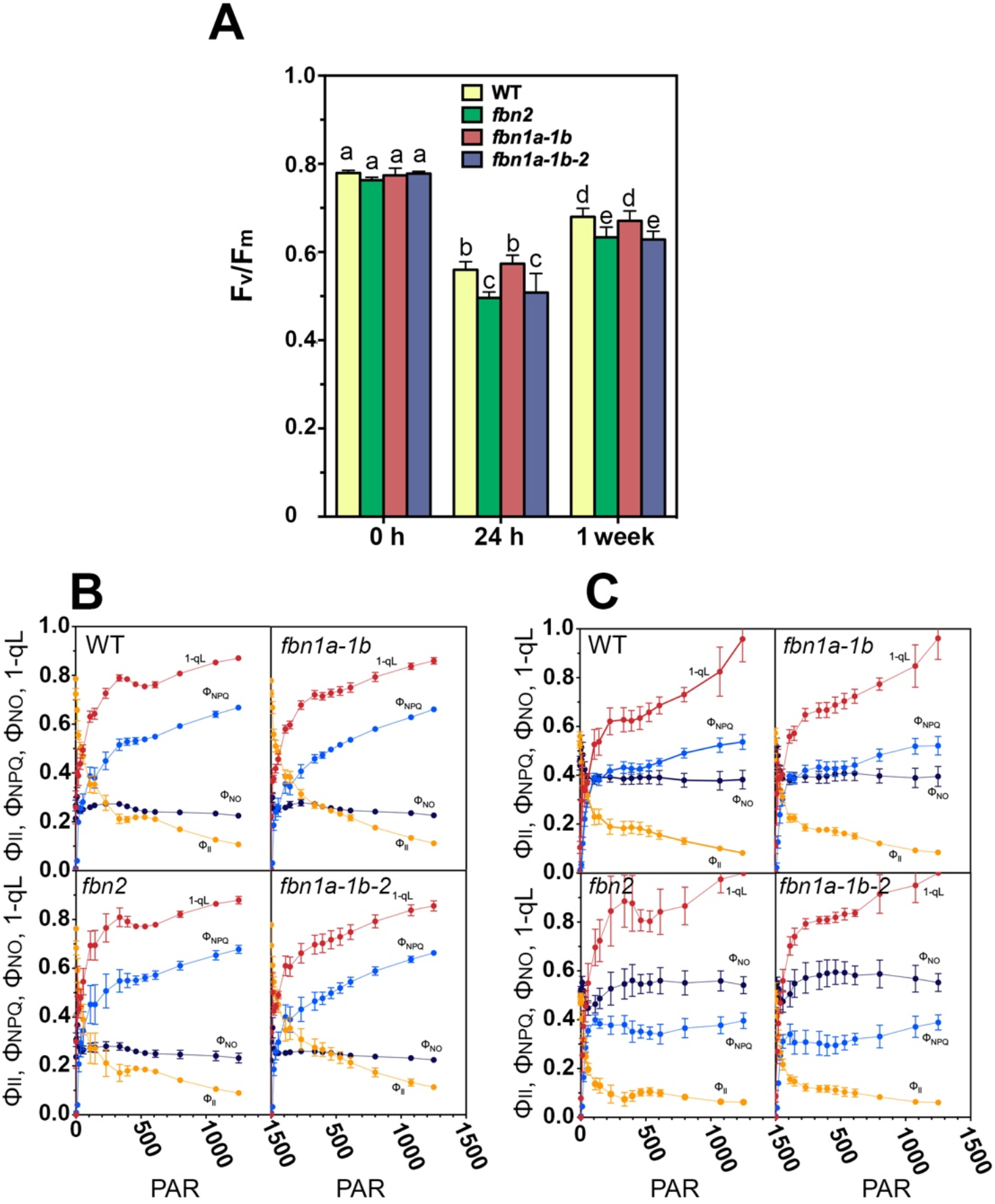
Photosynthetic parameters of *fbn* plants. The plants were sown in soil, cultivated under normal conditions for 3 weeks, and stressed for 1 week with a combination of moderate high light and low temperature as described in Figure 2. (A) Maximum PSII quantum yield (F_v_/F_m_) was determined at 0 h, 24 h, and 1 week of stress. ANOVA was performed using Prism software version 6.0. Tukey’s HSD was used as the post-hoc test. Significance is denoted if p<0.01, and mean values varying significantly at the 1% level are marked with different letters. (B) Different photosynthetic parameters were determined at increasing light intensities using an IMAGING-PAM fluorimeter in plants at time 0 or (C) 24 h of stress: Φ_II_ (effective PSII quantum yield), Φ_NPQ_ (quantum yield of regulated energy dissipation), Φ_NO_ (quantum yield of nonregulated energy dissipation), and 1-qL (percentage of primary acceptor of PSII reduced). Values are presented as the mean and standard deviation (SD) of six biological replicates. PAR = photosynthetically active radiance (μmoles m^−2^ s^−1^.

### Screening for proteins interacting with FBN2

The above results indicate that FBN2 played a more relevant role than FBN1a and FBN1b in protecting the photosynthetic apparatus against abiotic stresses. To characterize the mechanism of action through which this FBN exerts this protection, we searched for proteins that interact with FBN2. We performed co-immunoprecipitation (Co-IP) analysis using specific polyclonal antibody against the full-length FBN2 protein.

As shown above, FBN2 exhibited dual localization: stromal and in association with membranes. Thus, we decided to immunoprecipitate FBN2 in both fractions. In the case of the FBN2 population associated with PGs, precipitation of this protein could co-purify other proteins associated with these organelles but that do not interact with FBN2. To avoid this artefact, after separation of the soluble and membrane fractions, we solubilized the FBN2 from PGs by incubating samples with 0.01% (v/v) Triton X-100, the concentration of this non-ionic detergent that solubilizes FBN2 from plastoglobules (Figure S2) prior to Co-IP. We checked that the anti-FBN2 antibody does not recognize the FBN1a or FBN1b proteins by immunoblotting under native and denaturing conditions (Figure S3). Finally, a parallel analysis was performed using *fbn2* plants to identify proteins that co-immunoprecipitated in a non-specific manner. Proteins obtained by Co-IP were identified by mass spectrometry, and those also found using *fbn2* extracts were removed (mainly highly abundant proteins from the photosystems and light-harvesting complexes). We performed three independent Co-IP experiments and the proteins found in all replications are listed in Table 1 (FBN2 associated with PGs). In the case of the soluble population of FBN2, only isoforms 1 and 2 of fructose bisphosphate aldolase (FBA) were found with Co-IP. These isoforms have been described as proteins associated with PGs (Vidi et al., 2006; Ytterberg et al., 2006) but are not considered as part of the core proteome (Lundquist et al., 2012). The rest of the proteins that were identified seemed to interact exclusively with the FBN2 population associated with PGs. Some of those proteins were described previously as components of the core proteome of PGs (Lundquist et al., 2012) and are highlighted in bold in Table 1. Interestingly, the ABC1-type kinases, the second most abundant type of protein in PGs, were not found in these analyses, indicating that the identified proteins bind to FBN2 and were not mere contaminants of PG-associated proteins. Finally, other proteins have not been previously described as being associated with PGs, such as isoform 2 of the ferredoxin-NADP[H] oxidoreductase (FNR2), which forms a complex with Tic62 (also found by Co-IP) (Benz et al., 2009), or the APE1 protein, involved in adaptation of the plant, specifically the thylakoid membranes, to fluctuating environmental conditions (Walters et al., 2003).

**Table 1.** Proteins that co-immunoprecipitate with FBN2 Proteins previously associated with the plastoglobules are in bold. aDescribed by Lunquist et al. (2012). bDescribed by Vidi et al. (2006) and Ytterberg et al. (2006).

| AGI code | Name |
| --- | --- |
| **At4g04020** | **Fibrillin 1a, FBN1a** |
| **At2g21330<sup>b</sup>** | **Fructose bisphosphate aldolase 1** |
| **At4g38970<sup>b</sup>** | **Fructose bisphosphate aldolase 2** |
| At3g45140 | Lipoxygenase 2, LOX2 |
| **At5g42650<sup>a</sup>** | **Allene oxide synthase, AOS** |
| At5g38660 | Acclimation of photosynthesis to environment, APE1 |
| At1g20020 | Ferredoxin-NADP[H]-oxidoreductase 2, FNR2 |
| At3g18890 | NAD(P)-binding Rossmann-fold superfamily protein, Tic62 |
| **At4g19170<sup>a</sup>** | **9-cis-epoxycarotenoid dioxygenase, CCD4** |
| **At5g08740<sup>a</sup>** | **NAD(P)H dehydrogenase C1** |
| **At1g32220<sup>a</sup>** | **NAD(P)-binding Rossmann-fold superfamily protein** |
| At4g35250 | NAD(P)-binding Rossmann-fold superfamily protein |

### Interaction of FBN2 with FBN1a and other FBN2 polypeptides

One of the proteins that co-immunoprecipitated with FBN2 is FBN1a (Table 1). The possible interaction between these two proteins was confirmed by bimolecular fluorescence complementation (BiFC) during transient expression in *Nicotiana benthamiana* leaves. Figure 4 shows the *in vivo* interaction between FBN2 and FBN1a, with a dot-like pattern similar to that observed for SSIV-FBN1a (Gámez-Arjona et al., 2014b) or FBN1b-FBN1a (Gámez-Arjona et al., 2014a) interactions. We previously showed the head-to-tail interaction between two FBN1a or FBN1b polypeptides, suggesting that they can form homodimers, heterodimers, or oligomers *in vivo* (Gámez-Arjona et al., 2014a). Considering the homology between these proteins, we analyzed whether FBN2 could also bind to another FBN2 polypeptide. This interaction was confirmed *in vivo* when FBN2 was expressed transiently in *Nicotiana* leaves (Figure 4).

**Figure 4.**
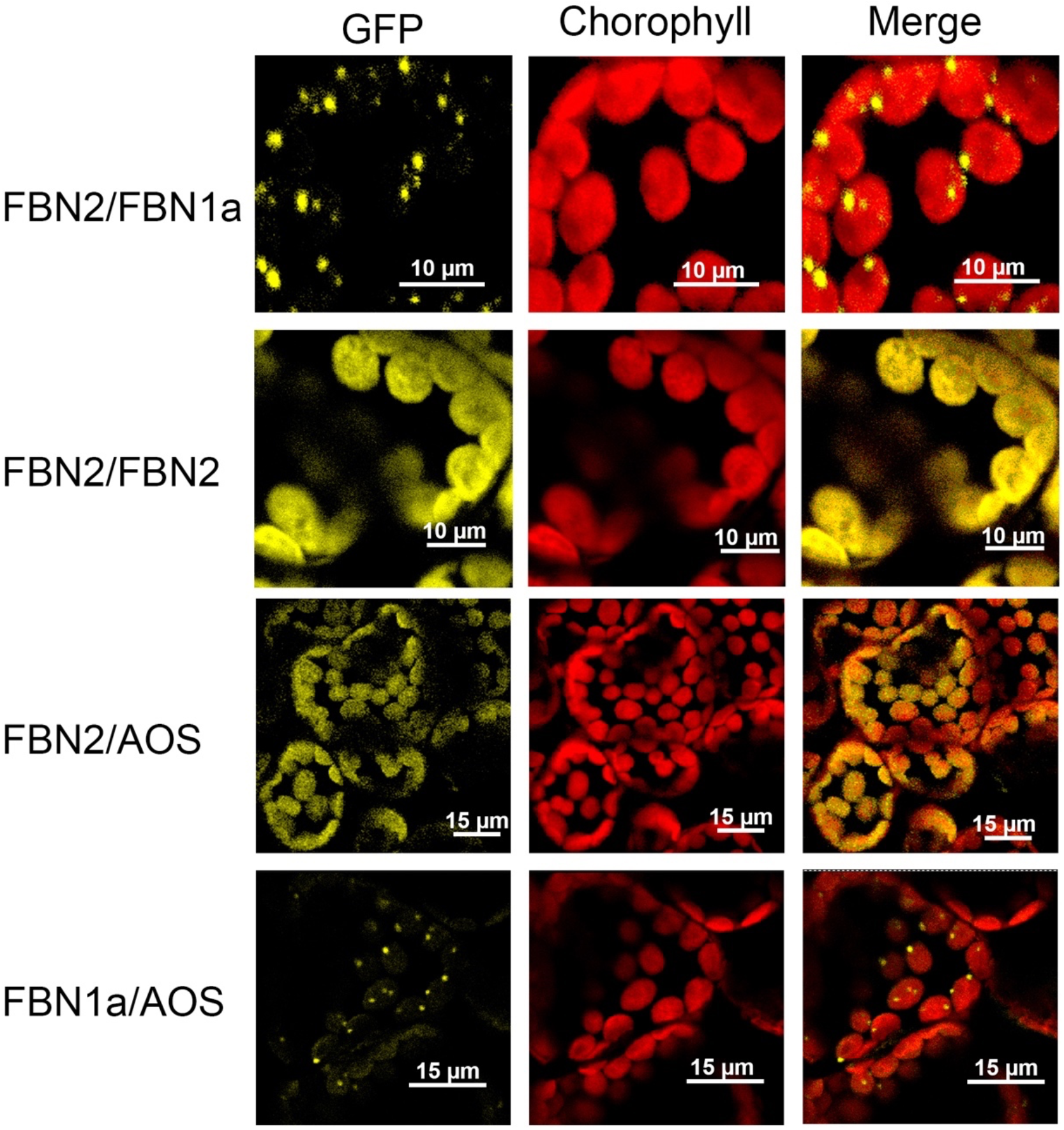
*In vivo* interaction between FBN2 or FBN1a and FBN2, FBN1a, or AOS polypeptides. cDNAs encoding the full-length FBN1a or FBN2 proteins were fused to the N-terminal half of YFP and co-transformed into *N. benthamiana* leaves together with cDNA encoding FBN2, FBN1a, or AOS fused to the C-terminal moiety of CFP. Images show the YFP/CFP (BiFC) fluorescence, the chlorophyll autofluorescence, and the merged images.

### FBN2 binds to allene oxide synthase

Co-IP analysis identified allene oxide synthase (AOS) as a protein that binds FBN2. This protein has been described as a component of the core PG proteome and catalyzes, together with lipoxygenase (LOX) and allene oxide cyclase (AOC), the first steps of the biosynthesis of JA in the chloroplast (Wasternack and Song, 2017). Lipoxygenase 2 (LOX2) was also identified in the Co-IP analysis (Table 1), and AOC was found in these analyses but was not included in the list because it did not appear in all three replications. Figure 4 shows that AOS can interact *in vivo* with FBN2, confirming the results obtained in the Co-IP analysis. In addition, we found that AOS is able to interact with FBN1a *in vivo* (Figure 4).

### Interaction between FBN2 and APE1

The Co-IP analysis identified some proteins that could *a priori* affect photosynthesis, such as FNR2 or NAD(P)H dehydrogenase C1, which is involved in reduction of the plastoquinone reservoir of plastoglobules (Eugeni Piller et al., 2011). Among these proteins, we selected APE1 for further characterization, as its absence alters chlorophyll fluorescence and acclimation responses to the growth light environment (Walters et al., 2003). First, we confirmed the interaction between FBN2 and APE1 in *N. benthamiana* leaves by BiFC. Figure 5D shows that both proteins interact *in vivo* when transiently expressed in *Nicotiana* leaves. A way of identifying genes encoding proteins involved in a particular process is the mining of co-expression data over a range of tissues and conditions (Usadel et al., 2009). We used the CORrelation NETworks tool CORNET3.0 (https://bioinformatics.psb.ugent.be/cornet/) to perform this analysis. We screened all deposited data on gene expression with no bias towards any particular type of condition or treatment, and we queried FBNs associated with PGs (FBN2, −4, −7a, 7b, and −8) and APE1. FBN1a and FBN1b were disregarded by CORNET3.0 because of their sequence similarity. Figure 5A shows that FBN2 and APE1 are the closest elements in the network that is formed, sharing two highly co-expressed genes. This result supports the idea that FBN2 and APE1 are involved in the same metabolic process. To analyze whether the elimination of APE1 could promote similar alterations in PSII performance as those observed in the mutants lacking FBN2, two independent T-DNA insertion knock-out mutants of APE1 were characterized. The same results were obtained with both mutants, and only those obtained with the SALK_123198 line are shown. The elimination of APE1 decreased the maximum quantum yield of PSII in plants subjected to a combined stress of moderate high light (600 μmols m^−2^ s^−1^) and cold (10°C) for 24 h (Figure 5C). This reduction was also observed in the *fbn2* and *fbn1a-1b-2* mutants (Figure 3A), but was more pronounced in the *ape1* mutant. The effective PSII quantum yield (Φ_II_) was also reduced in *ape1* plants under stress conditions for 24 hours in a similar way as the reduction observed in *fbn2* plants (Figure 5B). In contrast, the non-photochemical quenching (NPQ) along a curve of increasing light intensity was reduced in the stressed *fbn2* plants, but was similar to WT in *ape1* plants (Figure 5B).

**Figure 5.**
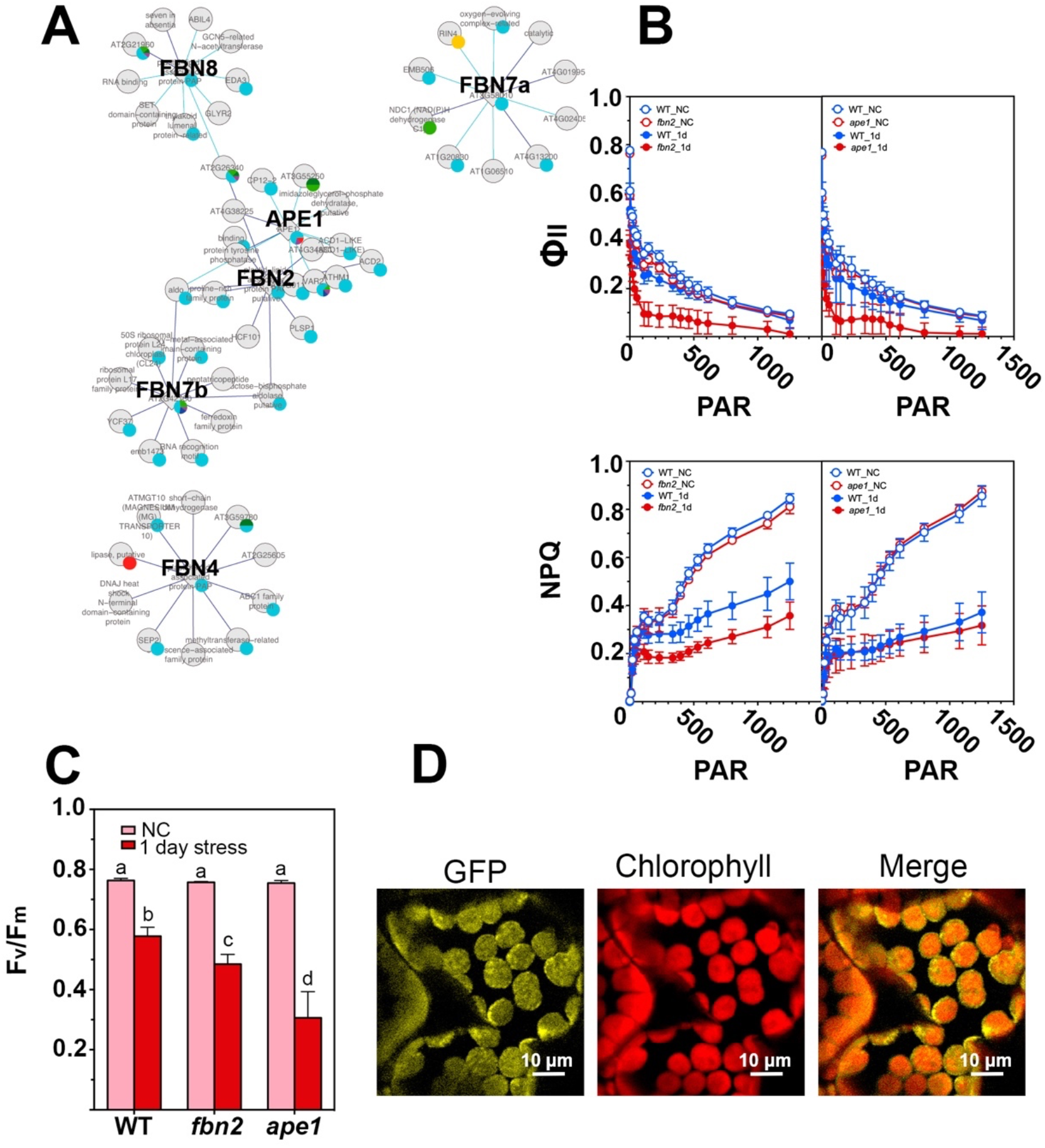
Relationship between FBN2 and APE1. (A) Coregulatory gene network of PG-associated FBNs and APE1 derived from condition-independent co-expression analysis using the CORrelation NETworks tool (CORNET3.0, https://bioinformatics.psb.ugent.be/cornet/). Only strong co-regulation of the top 10 genes with a Pearson correlation coefficient (corr. coeff) ≥ 0.8 (P≤0.05) are shown. Purple edge: corr. coeff ≥ 0.9. Cyan edge: corr. coeff ≥ 0.8. The relative size of a sector on the pie chart associated with each gene represents how much a protein is predicted to be localized to the compartment indicated (cyan: chloroplast, yellow: Golgi, red: extracellular, light green: mitochondria, dark green: nucleus, blue: cytosol, purple: plastid). (B) Effective PSII quantum yield (Φ_II_) and non-photochemical quenching (NPQ) at increasing light intensity of *fbn2*, *ape1,* and WT plants cultured under normal conditions at time 0 (open circles) or 24 hours (close circles) after a combined stress of moderate high light (600 μmols m^−2^ s^−1^) and low temperature (10°C). Values are presented as the mean and standard deviation (SD) of six biological replicates. PAR = photosynthetically active radiance (μmoles m^−2^ s^−1^). (C) Maximum PSII quantum yield (F_v_/F_m_) of the same plants analyzed in (B). ANOVA was performed using Prism software version 6.0. Tukey’s HSD was used as the post-hoc test. Significance was denoted if p<0.01, and mean values varying significantly at the 1% level are marked with different letters. (D) *In vivo* interaction between FBN2 and APE1. cDNA encoding the full-length FBN2 protein was fused to the N-terminal half of YFP and co-transformed into *N. benthamiana* leaves together with cDNA encoding APE1 fused to the C-terminal moiety of CFP. Images show the YFP/CFP (BiFC) fluorescence, the chlorophyll autofluorescence, and the merged images.

## DISCUSSION

FBNs have been proposed to be involved in central processes of plant physiology, such as plant responses to stress and plastid architectural development (Singh and McNellis, 2011). Some of these proteins have been suggested to be associated with PGs (Lundquist et al., 2012), though they could also be found in locations other than PGs. Thus, FBN4 was suggested to be present exclusively in association with PGs (van Wijk and Kessler, 2017), but it was also found to be a component of light-harvesting complex II (LHCII) subcomplex 3 (Galetskiy et al., 2008). FBN2 has been identified as part of the PG proteome core with a PG/stroma abundance ratio of 1,188 (Lundquist et al., 2012). However, the analysis of FBN2-GFP localization suggests that this protein is also present in the stroma or uniformly localized in the thylakoid membranes. Immunoblot analysis of plastidial soluble and membrane fractions indicated that FBN2 is distributed in similar proportions in these two fractions. We cannot establish that the membrane-associated population of FBN2 is localized exclusively in the PGs, and it could also be localized in other regions of the thylakoid membranes. The different methodology employed, mass spectrometry in Lundquist’s analysis and GFP fusion and immunoblot in this study, may explain the discordance between the two analyses. These results are also different from those obtained by the analysis of FBN1-GFP, which rendered a dot-like pattern of localization characteristic of PG-associated proteins (Gámez-Arjona et al., 2014b), and it is the first evidence pointing to different functions of FBN2 and FBN1s.

AOS was previously identified as part of the PG proteome core by Lundquist et al. (2012), and recruitment of the three plastidial enzymes (LOX, AOS, and AOC) in the synthesis of JA to the PGs of stressed plants has been shown (Lundquist et al., 2013). Recently, the existence of a chloroplast envelope complex comprising LOX2, AOS, and AOC was described (Pollmann et al., 2019). These data suggest the existence of two populations of enzymes, one localized at the chloroplast envelope and another associated with the PGs. Furthermore, according to Lundquist et al. (2013), these enzymes would be recruited to PGs under stress conditions. We have shown that LOX2 and AOS interact with FBN2 (and likely also with FBN1a), and that the elimination of FBN1s or FBN2 slows down the JA-mediated high light-induced accumulation of anthocyanins in the mutant plants. The suppression of this alteration when plants are treated with JA indicates that the elimination of FBN2 or FBN1s affects the synthesis of JA. An explanation of this effect is that FBN1s and FBN2 mediate the association of these enzymes with PGs, facilitating the flux of metabolites through them. The slowed accumulation of anthocyanins would be more noticeable in *fbn*-defective mutants subjected to abiotic stresses, a situation that increases the number of PGs in the cells (van Wijk and Kessler, 2017) and the amount of FBNs (Ytterberg et al., 2006). The recruitment of enzymes involved in JA synthesis to the PGs would be affected by the elimination of FBN2 or FBN1s. Nevertheless, we cannot discount that the interaction of these FBNs with the JA synthesis enzymes positively modulates the activity of these enzymes.

The function of these FBNs in JA synthesis do not seem to be redundant, as the same alteration was observed in the *fbn1a-1b*, *fbn2,* or *fbn1a-1b-2* mutants, indicating that FBN1s and FBN2 are necessary in this process and the elimination of any of them alters the synthesis of JA. In this regard, the interaction observed between FBN2 and FBN1a is interesting. We previously showed that FBN1a and FBN1b can form homopolymer or heteropolymer *in vivo* by interacting with each other via a head-to-tail mechanism (Gámez-Arjona et al., 2014a). These data suggest that these three FBNs may form a net on the surface of PGs, and we could speculate that this net is necessary for the recruitment of the enzymes involved in JA synthesis to PGs, so that the elimination of FBN1s or FBN2 would alter JA synthesis in stressed plants.

Other alterations are specific to *fbn2* mutants. Thus, the elimination of FBN2 affects PSII performance in light/cold-stressed plants (Figure 3). These plants have a lower maximum quantum yield of PSII (F_v_/F_m_), indicating more stress-mediated PSII lesions in *fbn2* mutants than in WT or *fbn1a-1b* plants. The *fbn2* mutants also demonstrate a lower effective photochemical PSII quantum yield (Φ_II_) and lower capacity to dissipate excess energy in a regulated manner (Φ_NPQ_), with the consequential increase in non-regulated dissipation of this excess energy (Φ_NO_). These alterations are not observed in *fbn1* plants (Figure 3). A clear relationship does not exist between FBN2 and these parameters, and these alterations should be analyzed in light of the Co-IP results. In addition to the enzymes involved in JA synthesis, the Co-IP analysis indicates that other proteins interact with FBN2 *in vivo* (Table 1), and these interactions could explain the observed modifications. In this regard, the identification of APE1 as an FBN2-binding protein is interesting. The dysfunction of APE1 in *fbn2* mutants could explain the decrease in the maximum and effective PSII quantum yields observed in these mutants. However, the decrease in the NPQ observed in *fbn2* was not found in *ape1* (Figure 5B), suggesting that other FBN2-interacting protein/s would be responsible for this alteration. APE1 is described as a thylakoid protein, with one transmembrane domain (https://www.uniprot.org/uniprot/Q2HIR7), and has not been found in any of the proteomic analyses of PGs (3-5). Our analyses show that FBN2 is not restricted to PGs and is also found in the stroma. Further studies will be necessary to ascertain if it is also located in other domains of the thylakoid membranes.

The Co-IP analysis suggests that the membrane-associated FBN2 population is a more active form than the soluble population, as just two proteins were found to bind the soluble FBN2 population, the most abundant plastidial isoforms of FBA in leaves; the other plastidial isoform is mainly expressed in roots (Lu et al., 2012). FBA catalyzes the condensation of fructose-1,6-biphosphate and the condensation of sedoheptulose-1,7-biphosphate in the Calvin-Benson cycle in plastids. Ytterberg et al. (2006) found these proteins in an analysis of the PG proteome and suggested that, as no other enzymes from the Calvin-Benson cycle was found, these aldolases made an unknown functional contribution to the metabolism/structure of PGs (Ytterberg et al., 2006). Further characterization will be necessary to understand the physiological meaning of the FBAs-FBN2 interactions and the mechanisms responsible for the partition of FBN2 into two compartments.

Our study suggests that the interaction with FBN2 is necessary for the correct functioning of different proteins, such as AOS or APE1. FBN2 could modulate or activate the function of these proteins. However, considering the broad spectrum of proteins identified in the Co-IP analysis, it is more likely that FBN2 acts as a dock that allows the correct positioning of different proteins, and its elimination would lead to the dysfunction of the interacting proteins.

## MATERIALS and METHODS

### Plant material and growth conditions

*Arabidopsis thaliana* plants were cultivated in a growth cabinet under different conditions: normal conditions, 16 h light/8 h darkness photoregimen, 110 μmol m^−2^ s^−1^ light intensity, 22°C day/20°C night temperature, and 70% humidity; high light stress, 16 h light/8 h darkness photoregimen, 600 μmol m^−2^ s^−1^ light intensity, 22°C day/20°C night temperature, and 70% humidity; cold stress, 16 h light/8 h darkness photoregimen, 110 μmol m^−2^ s^−1^ light intensity, 10°C temperature, and 70% humidity; and cold and high light stress, 16 h light/8 h darkness photoregimen, 600 μmol m^−2^ s^−1^ light intensity, 10°C temperature, and 70% humidity. The double mutant *fbn1a-1b* was described previously (Gámez-Arjona et al., 2014b). The *fbn2* (SALK_124590) mutant was obtained from the Salk T-DNA Mutants Collection (Alonso, 2003). T-DNA insertion mutants in APE1 are SALK_133182 from the Salk T-DNA Mutants Collection (Alonso, 2003) and 174C06 from the GABI-Kat Collection (Kleinboelting et al., 2012). The line carrying triple mutations, *fbn1a-1b-2*, was obtained by crossing and selecting homozygous triple mutant plants from the segregating F2 population using PCR-based genotyping. Table S1 describes all primers used.

### Plasmid construction

The full-length ORFs of FBN1a, FBN1b, FBN2, AOS, and APE1 were cloned (without the stop codon) into the pDONR207 entry vector (Invitrogen, http://www.lifetechnologies.com) using a BP reaction (Invitrogen). After sequence verification, the inserts were transferred into the binary vectors pXNGW(-*n*YFP) or pXCGW(-*c*CFP) (Yuan et al., 2013) for BiFC assays using LR Clonase II (Invitrogen). This resulted in translational fusion between the ORFs and the YFP/CFP moieties driven by the CaMV 35S promoter. FBN2 cDNA was also transferred to vector pGWB5, which allows the expression of translational fusion FBN2-GFP under control of the 35S promoter (Nakagawa et al., 2007).

### Transient expression in *N. benthamiana*

The *Agrobacterium*-mediated transient expression of the different genes in *N. benthamiana* leaves was carried out as described by Gámez-Arjona et al. (2014b).

### Confocal microscopy

A DM6000 confocal laser scanning microscope (Leica Microsystems, http://www.leica-microsystems.com) equipped with a 63x water-immersion objective was used to examine protein localization by GFP fusion or protein-protein interaction in BiFC assays involving *N. benthamiana* mesophyll cells. GFP or YFP/CFP expression and chlorophyll autofluorescence were imaged by excitation with a 488 nm argon laser and detection at 500–525 nm and 630–690 nm, respectively.

### Immunoblot analysis

Proteins were transferred from an SDS-polyacrylamide gel to a polyvinylidene difluoride (PVDF) membrane by electroblotting using a Trans-Blot SD transfer cell (Bio-Rad, www.bio-rad.com) according to the manufacturer’s instructions. Blots were probed with rabbit anti-FBN2 (GenScript, www.genscript.com at 1:5000 dilution, rabbit anti-glutamine synthetase (GS; Agrisera) at 1:5000 dilution, or chicken anti-psbA (Agrisera) at 1:8000 dilution, followed by horseradish peroxidase-conjugated goat anti-rabbit IgG serum (Bio-Rad) at 1:25,000 dilution for anti-FBN2 and anti-GS, or conjugated goat anti-chicken IgG serum (Agrisera) at 1:10,000 dilution for anti-psbA. The hybridization signal was detected using WesternBright Quantum (Advansta). The chemiluminescence was visualized using a Chemidoc Imaging System (Bio-Rad) running Quantity One software (Bio-Rad).

### Co-immunoprecipitation analysis

Chloroplast isolation and thylakoid membrane purification were performed following the procedure described by Koochak et al. (2019) from 30 g of leaves from plants cultivated under normal growth conditions. An aliquot of thylakoid membranes equivalent to 250 μg of protein was resuspended in 500 μl (final volume) of 100 mM phosphate buffer (pH 7) and supplemented with Triton X-100 at a final concentration of 0.01% (v/v) to extract FBN2 from the membrane fraction. The sample was incubated at 4°C for 30 min and then centrifuged at 100,000 × g for 1 hour at 4°C. The pellet was discarded and the supernatant employed for Co-IP analysis using polyclonal specific antibody against the whole FBN2 protein (produced by GENSCRIPT) and Dynabeads Co-Immunoprecipitation Kit (Life Technologies, http://lifetechnologies.com) following the manufacturer’s instructions.

### Protein sample preparation and LC-MS/MS analysis

Protein treatment and analysis was performed as described by Vowinckel et al. (2014), with some modifications. After Co-IP, protein samples were precipitated with acetone and the pellet resuspended in a 0.2% solution of RapiGest (Waters, www.waters.com) in 0.05 M ammonium bicarbonate. Dithiothreitol (DTT) at a final concentration of 5 mM was added and the samples incubated for 30 min at 60°C. Finally, iodoacetamide (IAA) was added at a final concentration of 10 mM and the samples incubated for 30 min at room temperature in the dark. Digestion was performed by incubating the solution with trypsin at a ratio of 1:40 (trypsin:protein) overnight at 37°C. After digestion, the equivalent of 1 μg of protein was analyzed on a Tandem Quadrupole Time-of-Flight mass spectrometer (AB/Sciex TripleTOF5600 Plus) coupled with a Nanospray III Ion Source (AB/Sciex) and nano-HPLC (EKsigent Ultra 2D). Peptide separation was carried out by removing impurities on an isocratic pre-column (C18 PepMap100 column NAN75-15-03-C18-PM, Thermo Fisher Scientific) using 0.1% formic acid and 5% (v/v) acetonitrile as the solvent at a flow rate of 3 μl min^−1^ for 10 min. Peptides were then eluted onto the analytical column using the incorporated electrospray emitter (New Objective PicoFrit column, 75 μm id × 250 mm, packed with Reprosil-PUR 3 μm) and separated on a linear gradient of 5-35% solvent B for 60 min at a flow rate of 250 nl min^−1^. Solvent A was 0.1% (v/v) formic acid, and solvent B was acetonitrile with 0.1% (v/v) formic acid. The ion source was operated with the following parameters: ISVF = 2600, GS1 = 20, CUR = 25. The data acquisition mode, using the DDA method, was set to obtain a high-resolution TOF-MS scan over a mass range of 400-1250 m/z, followed by MS/MS scans of 50 ion candidates per cycle (scan mass range 230-1500 m/z) with dynamic background subtraction, operating the instrument in high-sensitivity mode. The ion accumulation time was set to 250 ms (MS) or 65 ms (MS/MS).

The proteins were identified using the software ProteinPilot v5.0.1 (Sciex) with the method Paragon, and the *Arabidopsis* proteome database (www.Uniprot.org) in FASTA format fused to the contaminants of Sciex.

### Photosynthetic parameters

For analysis of chlorophyll a fluorescence, plants were acclimated to the dark for 30 min prior to measurements. Chlorophyll a fluorescence was monitored using the Walz MAXI-IMAGING-PAM chlorophyll fluorometer. A pulsed, blue measuring beam (1 Hz, intensity 4) was used to obtain the F_o_ value. Saturation pulses of 2700 μmol m^−2^ s^−1^ were applied for 0.8 s to determine F_m_ and F_m_’. The maximum quantum yield of PSII was calculated as (F_v_/F_m_) = (F_m_-F_o_)/F_m_. Φ_II_ (effective PSII quantum yield) was calculated as (F_m_’-F’)/F_m_’, Φ_NPQ_ (quantum yield of regulated energy dissipation) as 1-Φ_II_ −1/(NPQ+1+qL(F_m_/F_o_-1), Φ_NO_ (quantum yield of non-regulated energy dissipation) as 1/(NPQ+1+qL(F_m_/F_o_-1)), and qL as (F_m_’-F)/(F_m_’-F_0_’) × F_0_’/F. The NPQ was determined as (F_m_-F’_m_)/F’_m_.

### Pigment determination

For anthocyanin determination, pigments were extracted by incubating the leaf samples in 1 ml of acidic (1% HCl) methanol overnight according to the method described by Rabino and Mancinelli (1986). The absorbance of the extracts, clarified by centrifugation, was measured at 530 and 657 nm, and the formula A_530nm_-0.2A_657nm_ was employed to determine the anthocyanin level.

## SUPPLEMENTAL MATERIAL

**Figure S1.** Photosynthetic parameters of *fbn2*, *fbn1a-fbn1b*, and *fbn1a-fn1b-fbn2* under high light or cold stress

**Figure S2.** Solubilization of FBN2 by Triton X-100 treatment

**Figure S3.** Specificity of anti-FBN2 antibody

**Table S1.** Oligonucleotides used

## ACKNOWLEDGMENTS

We thank Alicia Orea for technical assistance with the confocal microscopy, and Rocio Rodríguez and José María Personat for the LC-MS/MS analysis.

